# Perceptual history acts in world-centred coordinates

**DOI:** 10.1101/2021.02.18.431805

**Authors:** Kyriaki Mikellidou, Guido Marco Cicchini, David C. Burr

## Abstract

Serial dependence effects have been observed using a variety of stimuli and tasks, revealing that the recent past can bias current percepts, leading to increased similarity between two. The aim of this study is to determine whether this temporal integration occurs in egocentric or allocentric coordinates. We ask participants to perform an orientation reproduction task using grating stimuli while the head is kept at a fixed position throughout the whole session or while alternating position from one trial to the next, from left (−20°) to right (+20°), putting the egocentric and allocentric cues in conflict. Under these conditions, allocentric cues prevail.

## Introduction

Perception depends not only on the stimuli impinging on our senses but is strongly conditioned by expectations and past perceptual experience. Many perceptual properties – such as orientation, numerosity and face perception – are systematically biased towards the recent perceptual experience (Cicchini et al., 2014; Cicchini et al., 2017; Fischer & Whitney, 2014; Liberman et al., 2014). This effect, known as *serial dependence*, probably reflects an optimisation strategy, where perceptual systems take advantage of temporal redundancies (the relative stability of the world) to improve signal to noise ratios, and hence efficiency (Cicchini et al., 2018). Much evidence suggests that serial dependence acts directly within perceptual circuitry, at early stages of information processing (Cicchini et al., 2021), including monaural auditory circuits (Hao Tam Ho et al., 2019) and primary visual cortex (V1) (St. John-Saaltink et al., 2016). In this study we show that serial dependence for orientation judgments is spatially selective in external, not retinal coordinates, reinforcing the notion that it is driven by the temporal continuity of the external world.

## Methods

We measured serial dependence for orientation perception, with participants periodically tilting their heads from side to side between trials to dissociate retinotopic from spatiotopic representations.

### Stimuli and apparatus

Figure 1 illustrates the experimental setup and timeline. Each trial began with the fixation point whose colour signalled which side the observer should position their head. After a 2700 ms pause, sufficient to complete the head displacement (Mikellidou et al., 2016), a grating patch stimulus was presented centrally (spatial frequency 0.3 cpd, contrast 25%, 500 ms, 3.2°full-width half-height), followed by a mask (random noise filtered at 0.3 cpd, contrast 50%, 1000 ms). Observers reproduced the perceived orientation of the patch by setting the orientation of a mouse-controlled virtual line marked at its ends by two small circles (diameter 0.2°). Participants confirmed their choice with the space bar and the reproduction cursor disappeared.

**Figure 1.**
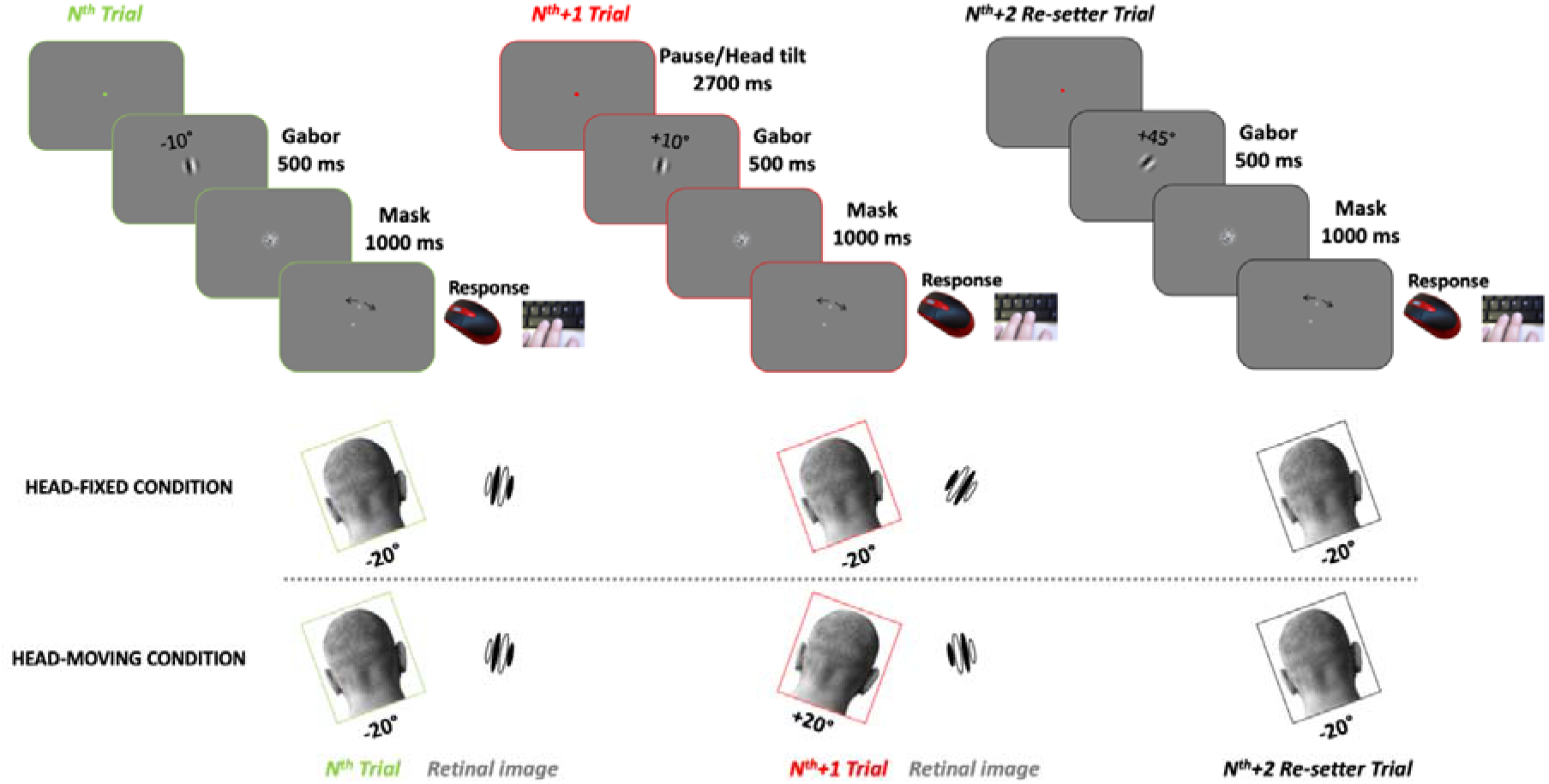
Timeline of experiment for the head fixed and head moving conditions, showing three consecutive trials. Trial sequence was designed so that the second trial of the triplet contained an orientation change of 20 degrees (in this example 20 degrees CW). This is paired with a change in head position of the same direction but twice in amplitude (40 degrees CW). The overall result is that the retinal image (shown in the inset only for the crucial trials) undergoes a rotation in the opposite direction (20 degrees CCW).

In a typical *head-fixed paradigm* (Figure 1 - top rows), egocentric and allocentric coordinate frames are the same. To dissociate between the two, we asked observers to alternate the position of their head from the left plate (approx. −20° from vertical) to the right plate (approx. +20° from vertical) from one trial to another (bottom row). When the head changes position by 40° and the stimulus rotates 20° in same direction, the rotation in allocentric coordinates is +20°and in egocentric (retinotopic) coordinates −20°, equal and opposite to that in allocentric coordinates. These trials were crucial to distinguish between the coordinate frames of the effects.

We concatenated triplets of trials in which the first two stimuli follow the above rule, and the third stimulus acts as a re-set, with its orientation chosen either 35 or 55° away from the preceding stimuli. Only data from the crucial trials were isolated and analysed. As orientation reproduction may exhibit biases due to attraction towards diagonals (or repulsion from cardinals (de Gardelle et al., 2010; Jastrow, 1892; Taylor & Bays, 2018), in order to pool data across orientation we first removed subjective biases at any orientation by subtracting the average response to that orientation in non-critical trials where there was an inter-trial difference exceeding 35°. We then estimated the weight of the previous stimulus by dividing the specific error to a given trial by the orientation difference with the previous trial. To estimate the egocentric and allocentric component of serial dependence we collected on each participant two sessions in fixed head condition and two in alternating condition (48 trials in each session-96 in total for each condition).

Stimuli were generated under MATLAB version 7.6 using Psychtoolbox routines^1^,^2^ and presented on a 23-inches LCD monitor (52° x 29°) with 1920 x 1080 resolution at a refresh rate of 60 Hz and mean luminance of 38 cd/m^2^. Observers viewed the stimuli binocularly from a distance of 57 cm.

### Participants

Sixteen participants were initially screened with the head-stationary condition. Those who showed positive serial dependence when the head was still were asked to complete the second part of the experiment, with the head alternating position from a left to a right head rest as described above. Eight participants (6 female (age range 23 – 40 years old) completed both conditions. All observers were naïve to the objective of the experiments, except authors K.M and G.M.C., and all had normal or corrected-to-normal vision. Experimental procedures were approved by the regional ethics committee (*Comitato Etico Pediatrico Regionale-Azienda Ospedaliero-Universitaria Meyer*, Florence) and are in line with the Declaration of Helsinki. All participants gave informed written consent.

## Results

Participants reproduced the orientation of briefly presented grating patches, while tilting their heads alternately between −20° (left) to +20° (right) between trials, as well as with their head stationary, as described above (see Fig 1A). Figure 2A shows data for all participants, plotting head-fixed against alternating-tilt conditions. If the effect depends on allocentric orientation of the stimulus, then tilting the head should make no difference to the magnitude of the effect, and the data should align with the positive (blue) diagonal. On the other hand, if the effect is egocentric (or retinotopic), the sign of the effect should invert (see methods), and the data should lie on the negative (green) diagonal. Clearly, they align with the positive diagonal. The likelihood that they do so is 1.56, compared with 0.0017 that they follow the egocentric prediction, giving a likelihood ratio (Bayes factor) of 917.

**Figure 2.**
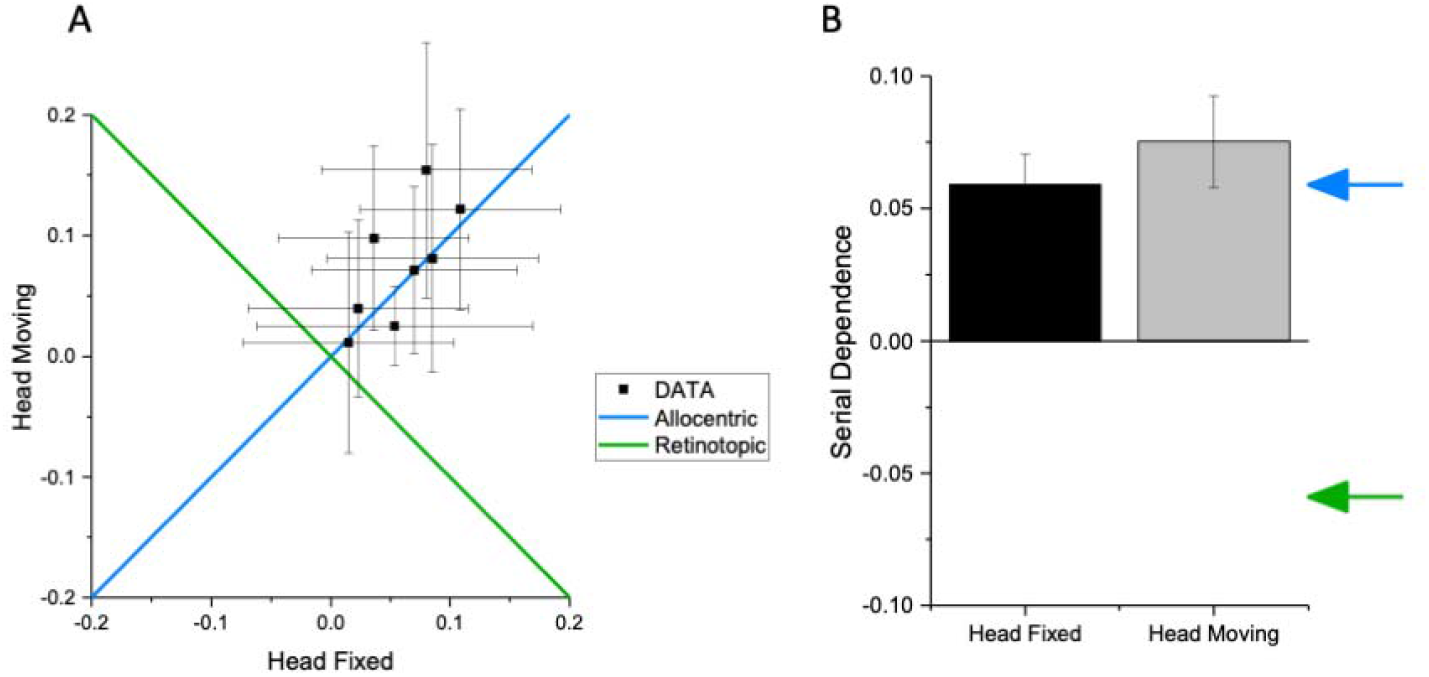
A. Individual data points for the head fixed and head moving positions clearly follow allocentric coordinates. B. Average bar plots for the head fixed (black) and head moving (grey) conditions, showing that serial dependence carries a strong allocentric component. Blue arrow indicates allocentric predictions; green arrow indicates retinotopic predictions.

Figure 2B shows the average results as bar plots, with the allocentric and egocentric predictions shown respectively by blue and green arrows. The average results clearly follow the allocentric predictions, with the effects measured in the head alternating condition statistically indistinguishable from those with head stationary (BF=0.65 (two tail), BF=0.174 (one tail)). The serial dependence was clearly allocentric, with no measurable effect corresponding to retinal orientation, which predicts a bias in the opposite direction.

## Discussion

Previous evidence for the coordinate system of serial dependence has been inconsistent. In their original study Fischer and Whitney (2014) observed allocentric tuning for orientation selectivity after saccadic displacement, while Collins (2019) reports clear retinotopic effects, using a saccade paradigm. One of the reasons for the difference may be that here we put retinotopic and allocentric signals in conflict, so the stronger one dominates. There may exist retinotopic effects, but these would seem to be overridden by more robust spatiotopic effects when one is pitted against the other, agreeing with a recent study on allocentric motion perception (Drissi-Daoudi et al., 2020).

If serial dependence serves to aid perceptual continuity, it would need to be allocentric. The external world tends to remain relatively constant over the short term, but this is not true of the retinal image: retinal position changes on each eye-movement, and retinal orientation changes when we tilt our heads. To exploit temporal perceptual redundancies, the system requires access to allocentric information, corresponding to external reality.

Nevertheless, the result is particularly interesting and perhaps unexpected in the light of evidence for serial dependence effects in early sensory cortex (St. John-Saaltink et al., 2016), which is commonly assumed to be retinotopically rather than allocentrically tuned. One could imagine several plausible explanations for this. One is that orientation signals of remembered stimuli may be reconverted back into retinotopic coordinates and projected back to V1 to interact with incoming signals. Indeed, feedback from higher areas may be one of the key mechanisms mediating serial dependence (Cicchini et al., 2021). Alternatively, even primary visual areas may display more allocentric properties than is commonly assumed. Indeed, there is good evidence for partial spatiotopy in primate V1 neurones during saccades (Trotter & Celebrini, 1999) and for allocentric space representation in early visual motion areas of humans (Crespi et al., 2011).

Our study provides clear evidence that serial dependence for orientation perception operates in allocentric coordinates, taking the inclination of the head into account. This is consistent with serial dependence reflecting the perceptual predictions, which remain constant over time in allocentric, but not egocentric coordinates.

## Acknowledgments

This project has received funding from the European Research Council (ERC) under the European Union’s Horizon 2020 research and innovation programme (Grant Agreement No 832813 – “GenPercept” to D.C.B.), and the Marie Skłodowska-Curie programme (grant agreement “Peripheality” No. 797603 to K.M.), from Flag-ERA JTC 2019 (grant “DOMINO” to G.M.C) and from the Italian Ministry of Education, University, and Research under the PRIN2017 program (grant number 2017SBCPZY—“Temporal context in perception: serial dependence and rhythmic oscillations” to D.C.B and G.M.C.).

